# Evolutionary stasis and homogeneous selection structure microbial communities in the deep subseafloor sedimentary biosphere

**DOI:** 10.1101/2025.11.18.689030

**Authors:** Tatsuhiko Hoshino, Atsuhiro Sakuma, Rei Kajitani, Hideyuki Doi, Takehiko Ito, Fumio Inagaki

## Abstract

The subseafloor biosphere, one of the Earth’s largest and most stable microbial habitats, mainly consists of energy-limited sediments where microbial life persists over geological timescales. However, the mechanisms governing microbial community assembly and evolution under condition of extreme energy limitation remains unclear. Here, we analysed a 296 m sediment core off the Shimokita Peninsula using amplicon sequencing, metagenomics, and genome-resolved analyses from 31 depth intervals spanning ∼480 kyr. The microbial community composition was governed primarily by homogeneous selection, consistent with persistent environmental uniformity during burial. Genome-resolved analyses of 224 high-quality metagenome-assembled genomes revealed highy conserved gene repertoires and uniformly low ratios of non-synonymous-to-synonymous substitutions, suggesting strong purifying selection with minimal genomic changes over this timescale. Our results indicate that evolutionary stasis is a pervasive feature of microbial life in deep subseafloor sediments, and propose a conceptual framework linking long-term environmental stability to genomic preservation in the Earth’s most persistent biosphere.

## Introduction

Understanding the evolutionary and adaptive processes of microbial life in subseafloor sediments, where cells persist for hundreds of thousands of years under condition of extreme energy limitation and physical isolation^1–3^, remains a central challenge in evolutionary microbiology. While surface ecosystems evolve in response to dynamic environmental and ecological pressures, deeply buried microbial communities experience prolonged periods with minimal disturbances over geological timescales.

The total number of microbial cells living in subseafloor sediments is estimated at 2.9–5.4 × 10²⁹, representing 0.18–3.6% of the planet’s total biomass ^2,3^. Cell densities are high in surface sediments but decrease by several orders of magnitude with depth, reflecting drastically reduced substrate and energy availability in deeper horizons^4^. Therefore, microbial communities in subseafloor sediments persist under progressively stronger energy limitation during burial^5^. Consequently, microbial growth rates in subseafloor sediments are expected to be extremely slow. Compared with the doubling times of microbial cells in energy-rich surface environments, those in sediments are markedly longer. Based on amino acid racemisation models, microbes inhabiting ∼1-million-year (Myr)-old sediments are estimated to double on annual-to-decadal scales^6^. Another estimate from abyssal red clay in the North Pacific reports respiration rates of ∼10⁻³ fmol O₂ cell⁻¹ day⁻¹ at 30 m depth, corresponding to potential doubling times on the order of thousands to millions of years^7^. In open-ocean Pacific sediments, estimates based on sulphate fluxes and cell densities suggest per-cell sulphate respiration rates of 1.5–12 × 10⁻²⁰ mol cell⁻¹ year⁻¹, which are far too low to sustain active movement such as a rotation of flagella^8,9^. Under such conditions, microbial dispersal is minimal; micrometre-scale diffusion coefficients suggest a displacement of only ∼6 m over 1 Myr^9^. Thus, the cells are effectively trapped within the sediment matrix and remain in place over geological timescales^10^.

Under such condistions, subseafloor microbial communities are distinct from those in seawater or surface sediments^11–13^. Species richness decreases sharply with depth, and the communities are dominated by specialised lineages adapted for long-term burial. Globally, members of the phyla Atribacterota (formerly the JS1 lineage) and Chloroflexota phyla consistently rank among the most abundant bacterial groups in anoxic marine sediments worldwide ^11,13,14^. These bacteria possess metabolic features that enable survival in subseafloor sediments. For instance, Dehalococcoidia can obtain energy by dehalogenating recalcitrant halogenated organic compounds^15,16^. Members of the Atribacterota rely on fermentative and syntrophic metabolisms to produce from amino acids and sugars both hydrogen and acetate, which are subsequently utilised by other microorganisms^17^. These interactions are essential for the long-term survival in energy-limited anoxic sediments^18^.

Despite considerable advances in the understanding of the diversity and metabolic potential of subsurface microbial communities, their assembly mechanisms and the genomic adaptation of individual taxa to extreme energy limitations remain poorly understood. Specifically, our understanding of microbial evolution during burial over thousands to millions of years remains elusive, with only a few pioneering studies addressing this complex topic, which have provided insightful data that suggest evolutionary stasis in microbial life across different subsurface settings. For instance, a notable investigation of genomic evolution in anoxic sediments from Aarhus Bay analysed four taxonomic lineages and found that their ratios of non-synonymous-to-synonymous substitutions (pN/pS) remained consistently below 1 across depths, suggesting persistent of purifying selection in anaerobic microbial groups within shallow horizons (<2 m; <5 kyr)^19^. However, data from deeper sediments remain scarce. Another study on *Thalassospira* strains from 3–6 Myr-old oxidised sediments at shallow depths (3 and 6 m below the seafloor [mbsf]) indicated that genetic drift and pseudogene formation might be enhanced by reduced effective population sizes in confined environments^20^. However, subsequent re-examination of the same *Thalassospira* isolates showed that their genomic features, including pN/pS ratios, pseudogene loads, and genome sizes, closely resembled those of their surface relatives. These findings suggest persistent purifying selection and minimal genomic changes over millions of years, likely reflecting extremely slow population turnover and effective genome maintenance under conditions of long-term energy limitation^21^. This phenomenon has also been observed in eukaryotes. The basidiomycete fungus *Schizophyllum commune*, recovered from ∼2 kmbsf, showed little genomic divergence from surface populations despite prolonged burial, although this system involves spore-forming microorganisms rather than metabolically active vegetative cells^22^. Beyond marine sediments, comparative genomics of Desulforudis audaxviator from deep continental aquifers revealed exceptional genomic conservation across globally distributed populations, with >99% average nucleotide identity (ANI) and nearly identical gene content.²² Despite geographical isolation, this remarkable stability has been attributed to extremely slow population turnover and efficient DNA repair, supporting the potential for long-term genomic stasis in geologically distinct subsurface environments.^23^. Collectively, these studies suggest that extremely energy-limited subsurface habitats suppress evolutionary processes, with microbial populations allocating their limited metabolic resources to maintenance and repair rather than to diversification, thereby enabling long-term persistence over geological timescales. However, systematic, depth-resolved investigations within the anoxic deep subseafloor, which constitutes the vast majority of the Earth’s sedimentary biosphere, remain largely unexplored.

In this study, we analysed a sediment core drilled in 2006 off the Shimokita Peninsula, extending 296 mbsf and encompassing sediments up to 480 kyr old. Our dataset of 31 depth horizons provides an unprecedented resolution for simultaneously characterising both community assembly and genome evolution during progressive burial. To disentangle the ecological processes governing community assembly, we performed 16S rRNA gene amplicon sequencing coupled with phylogenetic null modeling^24,25^. In parallel, we applied shotgun metagenomic sequencing to infer functional potential, and assembed metagenome-assembled genomes (MAGs) followed by comparative genomic analyses. By integrating community-level and genome-resolved approaches, we provide a comprehensive understanding of how bacterial populations adapt to and evolve in the deep coastal sedimentary biosphere.

## Results

### Microbial community composition

The vertical profile of the microbial communities at Site C9001 off the Shimokita Peninsula represents a typical pattern of anoxic marine sediments (Fig. 1), characterised by the dominance of anaerobic lineages such as Atribacterota and Chloroflexota^11^. These phyla dominated across all depths (Fig. 1a), with Atribacterota reaching nearly 60% relative abundance at 78 mbsf, whereas Chloroflexota consistently remained above 10% relative abundance throughout the sediment column. The anoxia-associated phylum Aerophobota was also detected at all the depths. Although the number of archaeal reads obtained was lower than the number of bacterial reads, two archaeal groups stood out. Asgardarchaeota (including Lokiarchaeia) comprised approximately 10–20% of the community in sediments down to 59 mbsf before declining sharply in deeper strata, whereas members of Hadarchaeota were rare in shallow sediments but increased to a maximum of 22% relative abundance below 200 mbsf. Microbial alpha-diversity (i.e., taxonomic richness) in the sediment samples decreased markedly with depth (Fig. 1B). Approximately, 3,600 amplicon sequence variants (ASVs) were detected in the shallowest sediments, but richness declined steeply toward 150 mbsf and stabilised at approximately 500–1,000 ASVs below that depth. Shannon’s diversity index showed similar trend, levelling off at ∼4 in sediments deeper than 150 mbsf. Beta-diversity analyses revealed a gradual turnover in community composition from the shallow to deep layers (Fig. 1c).

**Fig. 1.**
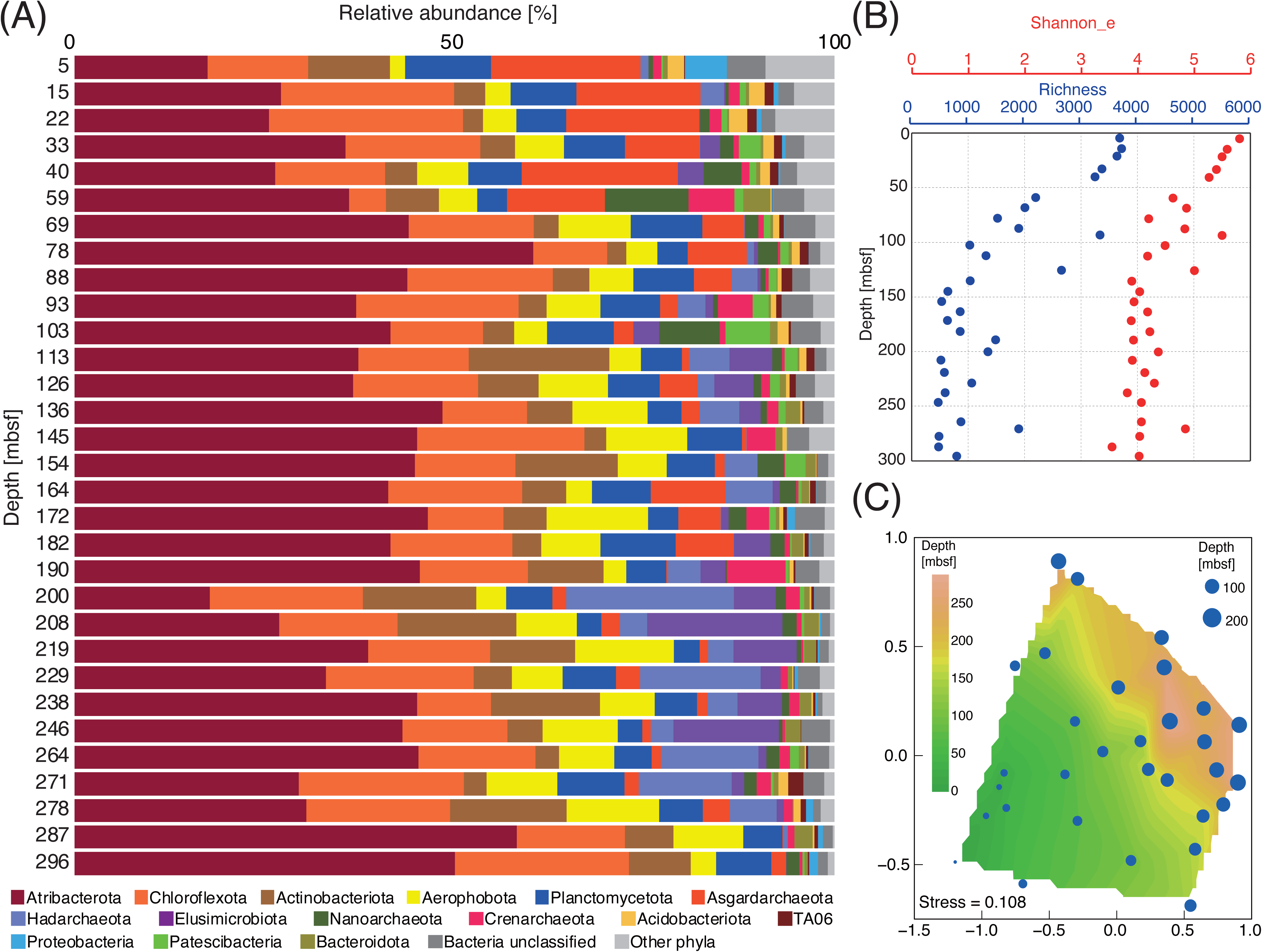
Microbial community composition. **a** Phylum-level community composition. Relative abundances of major microbial phyla across 31 sediment depth horizons (0–296 mbsf). **b** Shannon index based on the natural logarithm. Richness was defined as the number of ASVs in each sample. The ASV composition, rarefied to 200,000 sequences, was used for calculations. **c** Beta diversity of the microbial community composition. Non-metric multidimensional scaling ordination based on the Jaccard similarity index of rarefied ASV compositions. The sediment depths (mbsf) are shown on a contour map with a stress value of 0.108. *ASV* amplicon sequence variant, *mbsf* m below the seafloor, *Shannon_e* Shannon index.

### Community persistence across sediment depths

To explore the degree of connectivity and long-term persistence within the buried microbial community, we next examined whether specific taxa occurred consistently across all sediment depths. Due to the lack of active movement and cell division^6,7,9^, we infered that microbial assemblages formed in surface sediments were gradually buried and retained throughout the sediment column. If such energy-limited anoxic conditions persisted during burial, microbial taxa that are well adapted to them would be expected to persist from the shallow to deep layers. To identify persistent taxa, we extracted ASVs detected consistently across all sediment depths from the sequencing libraries. Of the 10,712 ASVs recovered across all samples,, 53 were shared among all depths. These persistent ASVs included Atribacterota (16 ASVs, 30%), Chloroflexota (13 ASVs, 25%), and Aerophobota (6 ASVs, 11%), with the remaining 18 ASVs belonging to various other taxa. Only one archaeal ASV, classified within Asgardarchaeota, was identified among the 53 persistent ASVs (Fig. 2a).

**Fig. 2.**
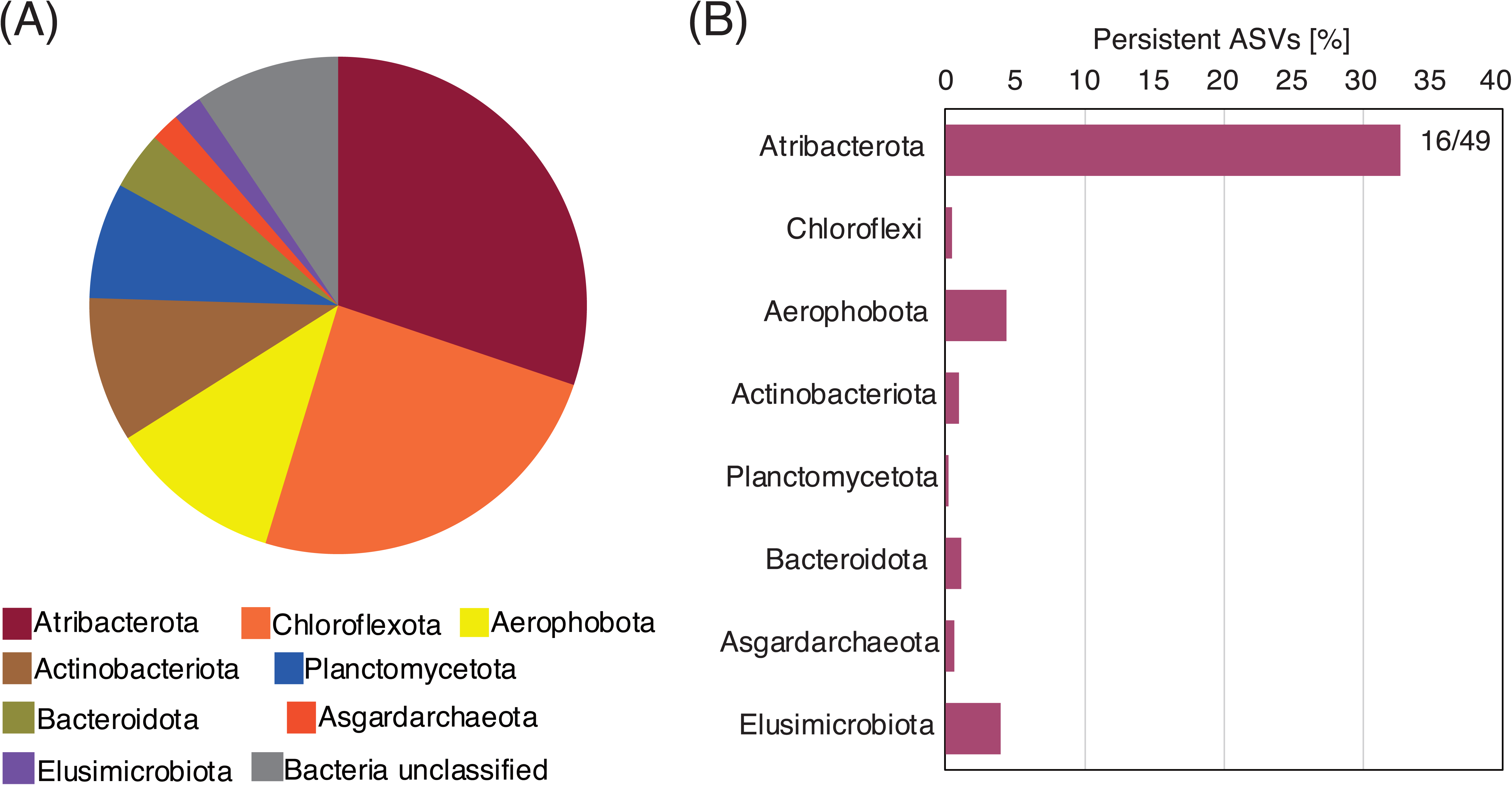
Taxonomic breakdown of ASVs shared across all sediment depths. Among the 10,712 ASVs detected across all sediment samples, 53 were consistently present at all depths. The pie chart illustrates the taxonomic affiliations (phylum level) of 53 ubiquitous ASVs. The bar chart on the right shows the number of shared ASVs (numerator) relative to the total number of ASVs assigned to each phylum (denominator), highlighting the lineages that include depth-persistent ASVs. *ASV* amplicon sequence variant.

The proportion of persistent ASVs varied notably among the phylogenetic groups. For Atribacterota, 16 out of 49 ASVs (33%) were consistently detected at all depths, in contrast to only 13 out of 2,946 Chloroflexi ASVs. Similarly low persistence was observed in other lineages, with only approximately 0.4–4% of ASVs detected at all depths, apart from Atribacterota. This pattern was further reflected in the depth profiles of ASV richness for each lineage. While the number of Atribacterota ASVs remained nearly constant across depths, ASV richness in other lineages declined with increasing depth, stabilising at ∼100 mbsf (a trend consistent with the data shown in Fig. 1B and Supplementary Fig. 1). These results suggest that in stratified sediments, where vertical microbial migration is unlikely, the observed community patterns are are due to selection processes that occurred during surface deposition rather than continuous adaptive changes during burial.

To test whether specific microbial lineages become increasingly dominant with increasing sediment depth as an adaptive response, we assessed the associations between depth and ASV relative abundance using Spearman’s rank correlation. Only a few ASVs exhibited a significant increase in their relative abundance with depth, whereas most ASVs either decreased or exhibited no significant trend. Together, these results indicate that the long-term persistence of the microbial community in deep sediments is not driven by adaptive dominance but instead reflects legacy effects established during initial burial. Consistent with this interpretation, deeper sediments showed neither a selective loss of taxa nor an adaptive increase in specific lineages, suggesting that community composition remains largely stable with depth. (Supplementary Table 1).

### Mechanisms of microbial community assembly

The processes shaping the microbial community structure in the deep subseafloor sedimentary biosphere are expected to differ fundamentally from those in other surface biospheres on Earth. Ecological processes underlying community assembly are generally classified into deterministic and stochastic categories. Deterministic processes include heterogeneous and homogeneous selection, whereas stochastic processes include dispersal limitation, homogenising dispersal, and drift^24,26^. To characterise the microbial community assembly in deep marine sediments at Site C9001, we applied the iCAMP pipeline^27^, which quantifies the relative influence of deterministic and stochastic ecological processes on community structure.

The results showed that the relative contributions of deterministic and stochastic processes in subseafloor sediments (46% and 54%, respectively) differed markedly from those in other biospheres, including air, ocean, and topsoil (Fig. 3A). Deterministic processes played a proportionally greater role in subseafloor sediments, primarily because of the exceptionally high contribution of homogeneous selection (42%), which was substantially higher than values reported for air (7.9%), ocean (7.6%), and topsoil (23.5%)^28^. The relative contribution of each ecological process varied across microbial lineages (Fig.3B). Homogeneous selection dominated the assembly of Atribacterota, accounting for nearly 70% of its total contribution, whereas dispersal limitation was the major process for most other phyla, typically exceeding 50% and reaching up to 70% in Chloroflexota. These lineage-specific differences indicate that Atribacterota are shaped primarily by consistent environmental filtering, whereas most other taxa are structured by spatial isolation in deep subseafloor sediments.

**Fig. 3.**
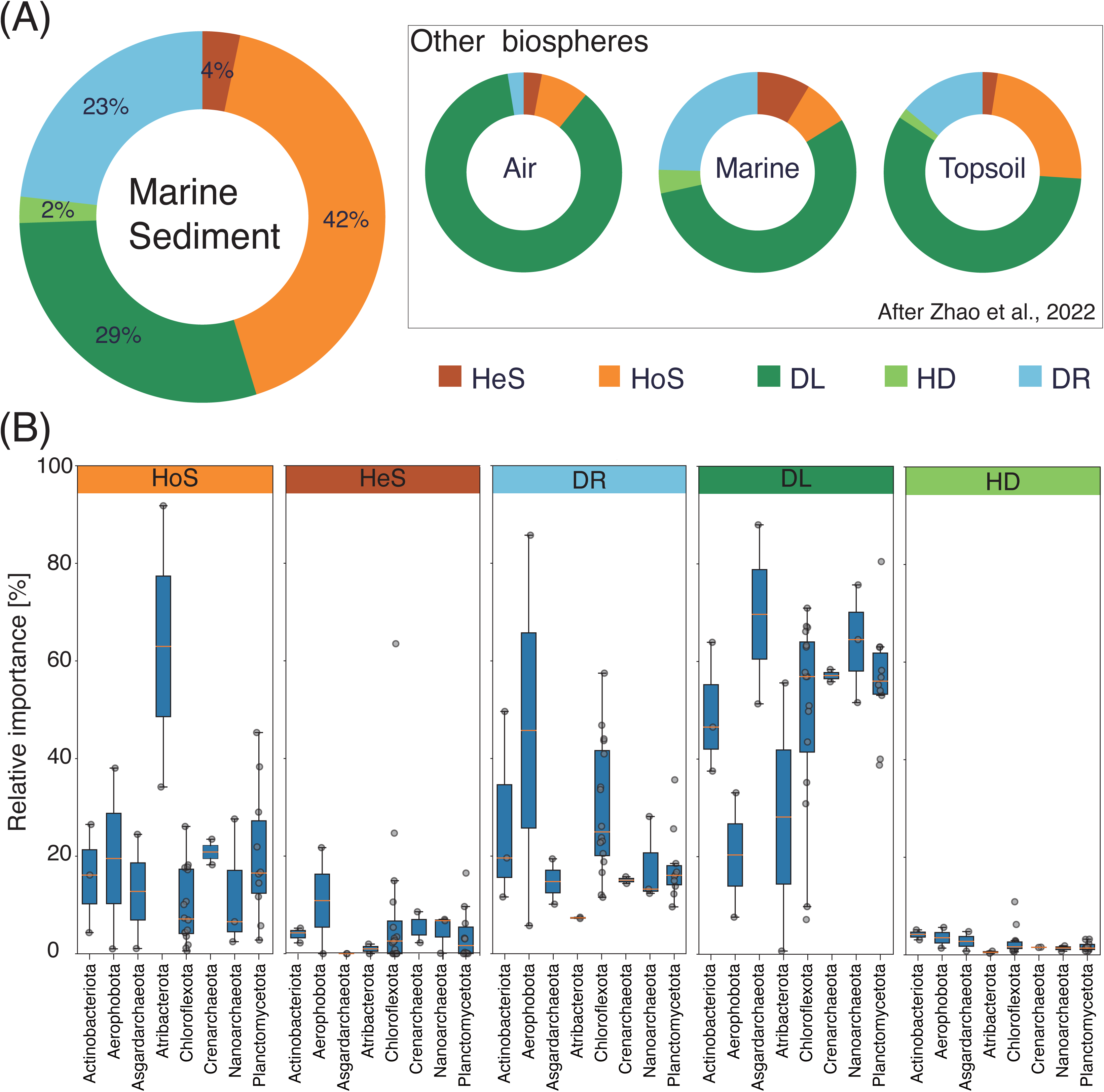
Ecological processes governing microbial community assembly in subseafloor sediments. Relative contributions of homogeneous selection (HoS), heterogeneous selection (HeS), dispersal limitation (DL), homogenising dispersal (HD), and drift (DR) were estimated using a phylogenetic null model framework. **a** Depth profiles showing process contributions across the sediment column. **b** Process contributions for individual microbial lineages.

### Phylogenetic Diversity and Distribution of High-Quality MAGs

A total of 3,993 genome bins were recovered from the subseafloor sediments, including 224 high-quality (HQ) and 1,017 medium-quality (MQ) genomes (Supplementary Fig. 2). High-quality MAGs consisted of 187 bacterial and 37 archaeal genomes, with Chloroflexota, Planctomycetota, and Atribacterota being the dominant bacterial phyla and Asgardarchaeota dominating the archaeal community. To resolve their evolutionary relationships, we constructed phylogenetic trees of 187 bacterial and 27 archaeal high-quality MAGs (Fig. 4). The bacterial tree revealed significant differences in the internal diversity of major phyla. For instance, the phylum Chloroflexota exhibited high phylogenetic diversity, with its MAGs branching deeply and spanning a wide area. In contrast, MAGs from the phylum Atribacterota formed a tightly clustered clade, indicating low phylogenetic diversity. These genomic observations were consistent with our 16S rRNA gene amplicon analysis, which also identified high diversity among Chloroflexota ASVs and notably low diversity among Atribacterota ASVs. Furthermore, the analyses of the MAG recovery depths showed that these key phyla were distributed across a broad depth range, rather than stratified into specific horizons. A similar trend was observed in the archaeal phylogenetic tree (Fig.4b); members of the dominant Asgardarchaeota were also recovered from throughout the sediment column, suggesting that diverse bacterial and archaeal lineages coexisted across the investigated subseafloor environment rather than being restricted to specific depth horizons.

**Fig. 4.**
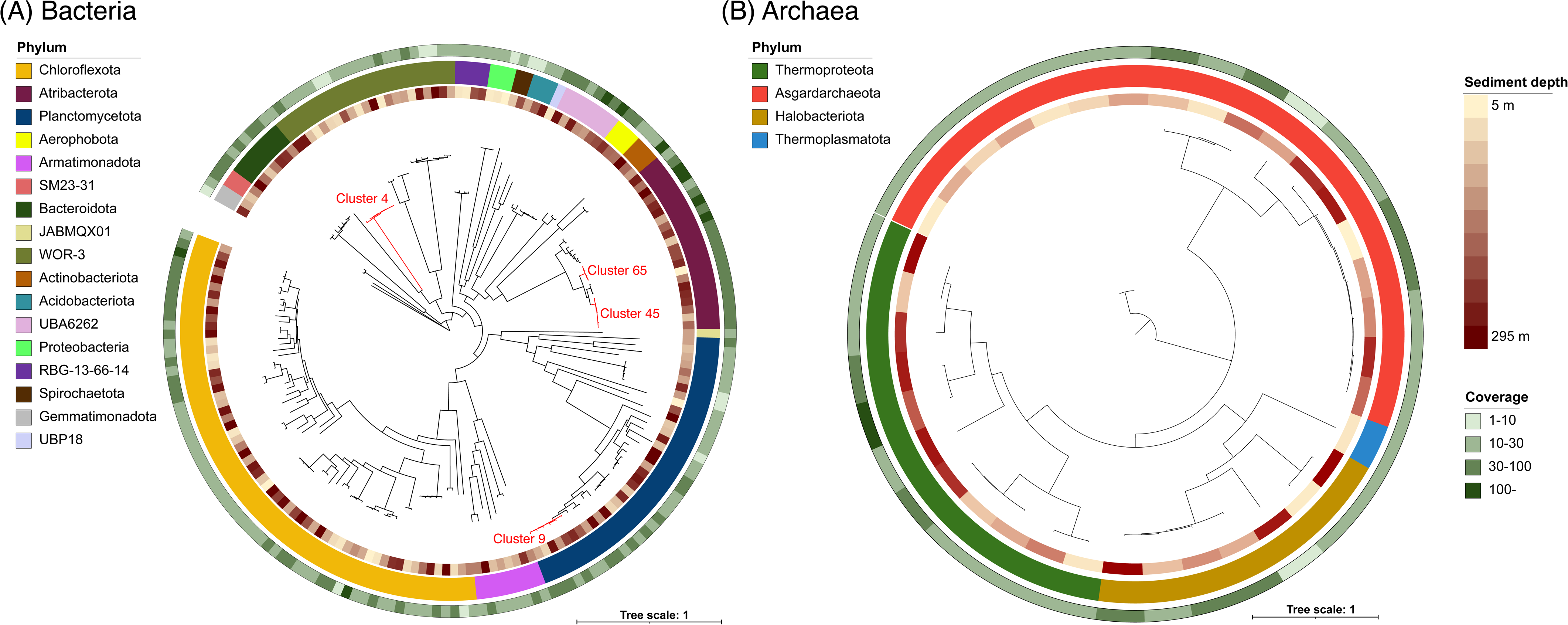
Phylogenetic trees constructed using the GTDB-Tk. The trees of (**a**) 197 bacterial and (**b**) 27 archaeal high-quality MAGs obtained in this study. The concentric rings surrounding each tree represent, from innermost to outermost, the sediment depth from which the MAG was recovered (5 m to 295 m), the phylum-level classification, and the average MAG coverage. In the bacterial tree (**a**), clusters marked in red text indicate groups of six or more MAGs with an ANI>97%, which were selected to analyse recombination-to-mutation ratios. The scale bar indicates the expected number of substitutions per site. *ANI* average nucleotide identity, *GTDB-Tk* Genome Taxonomy Database Toolkit, *MAG* metagenome-assembled genome.

### Trends in genome size by depth

Because genome streamlining has been proposed as a possible adaptation to energy-limited environments, we examined whether microbial genome size and coding ratios change systematically with sediment burial depth. Community-level estimates based on MicrobeCensus^29^ indicated that the average genome size was ∼4.7 Mb throughout the ∼300 m-deep sediment column at Site C9001, without a consistent reduction with increasing depth (Fig. 5). Lineage-specific estimates derived from high-quality MAGs revealed that members of Atribacterota and Dehalococcoidia typically possessed relatively small genomes of approximately 1.5–2.5 Mb, whereas Lokiarchaeia exhibited larger genomes of approximately 4–5 Mb. Across all examined lineages, genome sizes spanned approximately 1–8 Mb but did not significantly vary with depth. The coding ratios of these MAGs varied somewhat among lineages and depths but showed no clear correlation with sediment burial depth (Supplementary Fig. 3). These results indicate that neither genome size nor coding density changed significantly during long-term sediment burial.

**Fig. 5.**
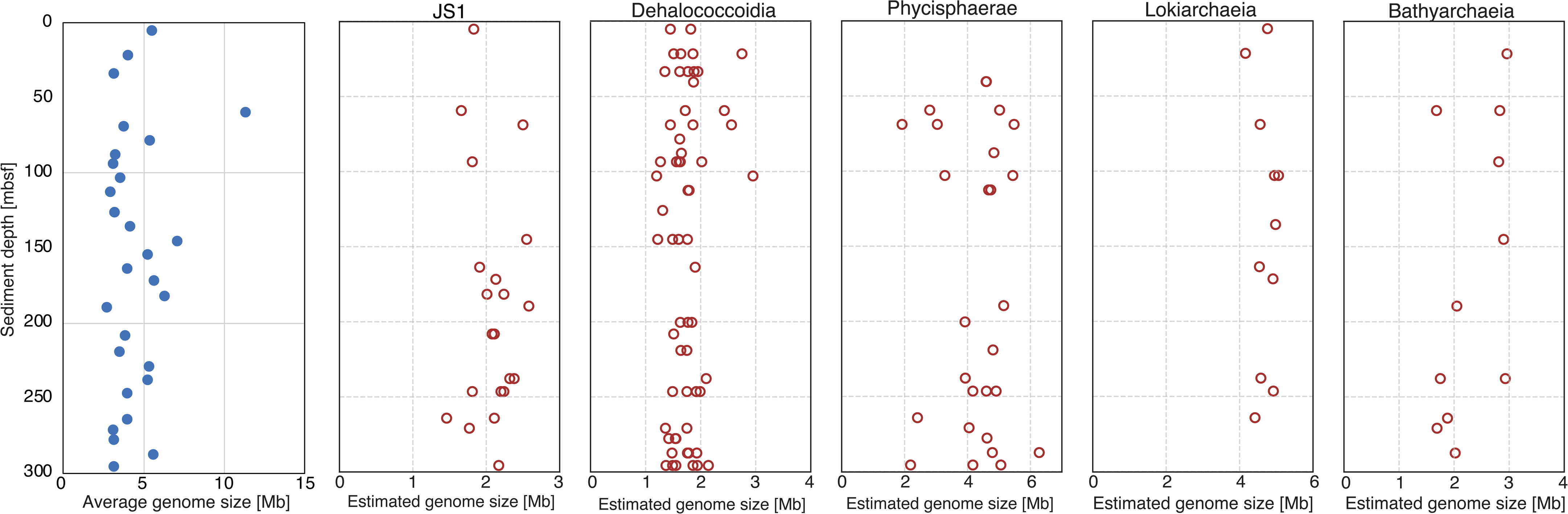
Genome size as a function of sediment depth. The left panel (closed blue circles) shows the average genome size of the entire microbial community in each sample, as estimated using MicrobeCensus. The other panels (open red circles) show the estimated genome sizes of individual MAGs from selected taxonomic groups: Atribacterota JS1 clade, Dehalococcoidia, Phycisphaerae, Lokiarchaeia, and Bathyarchaeia. The genome size for each MAG was estimated based on its total assembly length and completeness. *MAG* metagenome-assembled genome, *mbsf* m below the seafloor.

### Mutation rate and pN/pS ratios

To assess potential depth-related genomic variations, we compared mutation rates and pN/pS ratios between surface sediments and deeper horizons for four major bacterial lineages: Atribacterota, Chloroflexota, Aerophobota, and Actinomycetota. For each lineage, orthologous genes were identified and clustered based on ≥97% amino acid and nucleotide identity, enabling consistent comparisons of sequence variation across depths (Fig. 6). Across all four lineages, the distributions of both mutation rates and pN/pS ratios remained largely stable between the shallowest and deeper horizons. Mutation rates were consistently low, and the pN/pS ratios indicated strong purifying selection in all layers examined. In Atribacterota, the median pN/pS ratio was approximately 0.3 in both shallow and deep samples. Similar patterns were observed for Chloroflexota and Aerophobota, with pN/pS values generally ranging between 0.1 and 0.4 across depths. Actinomycetota exhibited somewhat broader distributions of pN/pS values, but these remained largely consistent with depth.

**Fig. 6.**
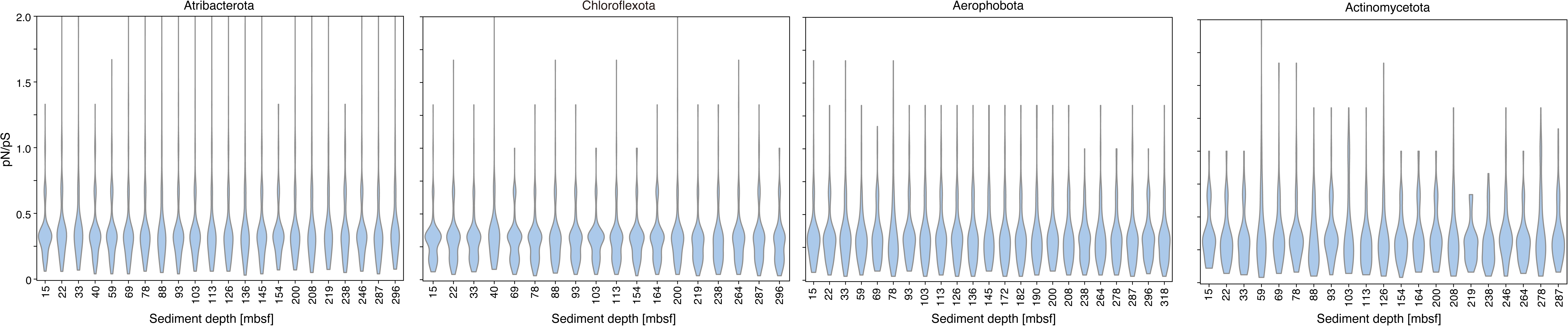
pN/pS ratios as a function of sediment depth. pN/pS ratios for representative microbial lineages across sediment depths at Site C9001 Each panel shows individual MAGs affiliated with major taxa (Atribacterota, Chloroflexota, Aerophobota, and Actinomycetota). pN/pS ratios were calculated from single-copy orthologous genes identified within each MAG. *MAG* metagenome-assembled genome, *mbsf* m below the seafloor, *pN/pS* non-synonymous-to-synonymous substitution.

To further test whether nucleotide substitutions accumulated with depth, we compared the sequence similarities of orthologous genes across all sediment horizons (Supplementary Fig. 4). For all four lineages, the average nucleotide identities of the orthologous genes remained uniformly high, without any systematic decrease from shallow to deeper horizons, indicating minimal accumulation of mutations with burial depth.

### Recombination-to-mutation ratios

We estimated the relative impact of homologous recombination on genome evolution across species-level MAG clusters (ANI ≥97%, shown in Fig. 4) by calculating recombination-to-mutation (r/m) ratios as the product of three parameters inferred by ClonalFrameML: the relative rate of recombination to mutation (R/θ), the mean length of imported DNA fragments (1/δ), and the nucleotide divergence of imports (ν; Table 1). Among the reference clusters from the surface or shallow subsurface environments, the r/m values varied widely (0.867–7.748). In contrast, subseafloor MAG clusters from this study showed a more moderate range (1.442–5.231). Both Atribacterota clusters exceeded their reference counterparts (2.841 and 5.231 vs. 1.079 and 1.816). The Planctomycetota cluster had an intermediate value of 3.244, whereas the WOR-3 cluster had a relatively low value of 1.442. Overall, the r/m estimates in subseafloor MAGs consistently fell within a moderate range (1–5). These r/m values contrast with the extremely low ratios (<1) reported for Thalassospira isolates from oxic subseafloor sediments^20^ and for Schizophyllum isolates recovered from >2 km-deep anoxic sediments^22^, indicating that the subseafloor populations analysed here exhibit comparatively higher r/m ratios. Owing to the extremely small biomass and slow growth rates in deep subseafloor sediments, such recombination events were most likely confined to near-surface sediments prior to or during early burial rather than occurring throughout progressive burial into deeper, energy-limited environments.

**Table 1.**
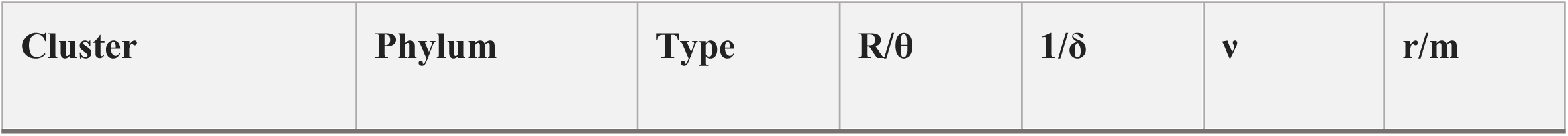
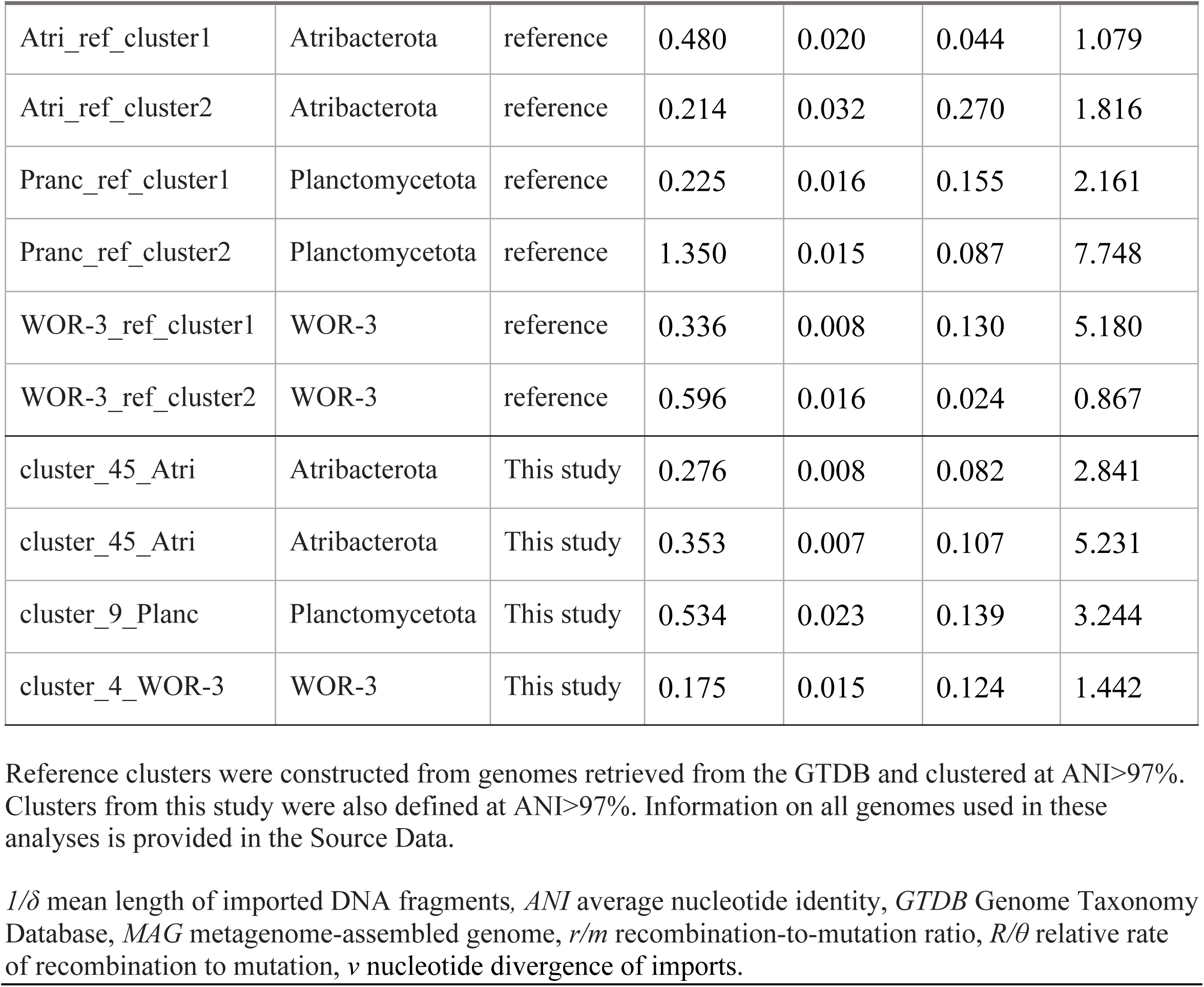
Parameters and estimates of r/m ratios in reference and subseafloor MAG clusters.

### Density of carbohydrate-active enzymes (CAZymes)

CAZymes are central to the degradation and recycling of complex organic matter in sediments^30,31^. Therefore, the depth-dependent distribution of CAZymes in the sediment column at Site C9001 may provide insight into how microbial lineages contribute to biogeochemical carbon cycling during long-term burial. We therefore evaluated the depth-dependent distribution of CAZyme gene densities in three taxonomic groups for which ≥ 15 high-quality MAGs were available, namely Atribacterota JS1 clade, Dehalococcoidia, and the Chloroflexota GIF9 clade, representing the only lineages with sufficient genomic replication for robust statistical comparison (Fig. 7, Supplementary Figs. 5 and 6). For Atribacterota and GIF9, no significant differences in CAZyme gene densities were observed across the sediment depth intervals (0–100, 100–200, and 200–300 mbsf) for any of the six CAZy categories (glycoside hydrolases, glycosyltransferases, carbohydrate esterases, auxiliary activities, carbohydrate-binding modules, and polysaccharide lyases), indicating a broadly consistent carbohydrate metabolic potential with burial depth. In contrast, Dehalococcoidia MAGs showed significant differences in glycoside hydrolase gene densities between 0–100 m and 100–200 m (p=0.0056), and between 100–200 m and 200–300 m (p=0.021). Carbohydrate esterase densities also differed significantly between 0–100 m and 100–200 m (p=0.0189), although no significant difference was observed between 100–200 and 200–300 m. These depth-associated patterns were detected only in Dehalococcoidia, in which significant differences were confined to a few CAZyme categories and depth pairs. Such limited variations do not constitute a consistent depth-dependent trend, supporting the overall stability of the carbohydrate metabolic potential throughout the sediment column.

**Fig. 7.**
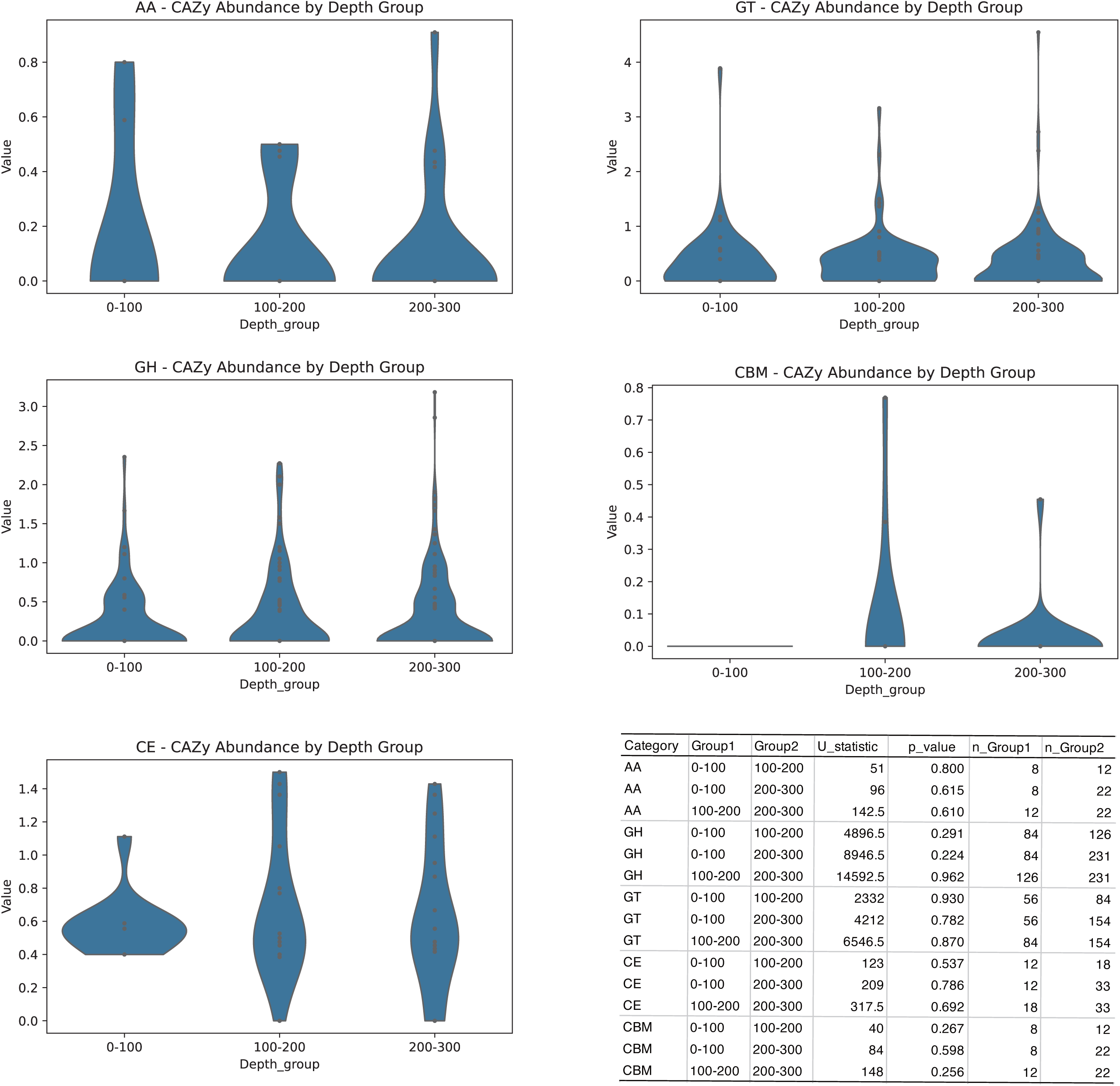
Depth-dependent variation in CAZyme gene density across MAGs affiliated with Atribacterota. Violin plots show the distribution of normalised CAZyme gene densities (genes per Mb) across three sediment depth intervals (0–100, 100–200, and 200–300 mbsf). CAZyme categories include auxiliary activities (AA), glycosyltransferases (GT), glycoside hydrolases (GH), carbohydrate-binding modules (CBM), and carbohydrate esterases (CE). No significant differences were observed across depths for any category (Mann–Whitney U test, p>0.2). *CAZyme* carbohydrate-active enzyme, *MAG* metagenome-assembled genome, *mbsf* m below the seafloor.

## Discussion

Among the processes shaping microbial community composition in the deep subseafloor sedimentary biosphere, homogeneous selection has emerged as the dominant process, highlighting the distinct environmental characteristics of this particular geomicrobiological setting relative to other environments on the Earth’s surface. Homogeneous selection refers to the assembly of similar microbial communities under relatively stable environmental conditions^26^. In the deep subseafloor, extreme and persistent energy limitation, combined with stable low temperatures, likely acts as a uniform environmental filter that selects for highly specialized metabolic strategies over geological timescales. The inherent environmental stability and chronic energy scarcity of the subseafloor sedimentary biosphere therefore make the dominance of homogeneous selection both plausible and consistent with current understanding of ecological dynamics in energy-limited subsurface ecosystems. The depth profiles of microbial diversity suggest that sediments 100 mbsf maintained greater environmental stability than shallower layers. When comparing the ecological processes driving community assembly between shallow (<100 mbsf) and deep (>100 mbsf) sediment bins, homogeneous selection accounted for a greater fraction in the deep bin (Supplementary Fig. 7). This pattern implies that deeper microbial communities are assembled more strongly by uniform environmental filtering than by stochastic processes, possibly because of subtle geochemical or mineralogical heterogeneities under otherwise geophysically stable sedimentary conditions. In contrast, dispersal limitation was the dominant process for most microbial lineages apart from Atribacterota (Fig. 3B). This pattern reflects the physical isolation of subseafloor sediments, which are largely closed systems with extremely limited cell migration between layers^9,19^. The pronounced role of dispersal limitation further suggests that microbial populations in subseafloor sediments have remained effectively segregated across sediment horizons, indicating that these communities have experienced long-term physical and ecological isolation with minimal opportunities for genetic exchange or recolonisation. Consistent with this observation, depth-resolved analysis showed that only 12 out of 10,712 ASVs increased in relative abundance with depth, whereas 1,948 ASVs significantly decreased; the vast majority showed no significant correlation with depth (Fig. 2, Supplementary Fig. 1, and Supplementary Table 1). The lack of depth-dependent dominance suggests that the adaptive expansion of particular lineages is rare in the deep sedimentary biosphere. This finding supports the view that persistent taxa primarily reflect legacy effects established at the time of burial rather than adaptive selection occurring during progressive burial in the deep subseafloor sedimentary biosphere. Therefore, deep subseafloor sediments therefore appear to select for specialised, well-adapted microbial taxa under stable, energy-limited conditions, while limiting both dispersal and large-scale ecological drift. This mode of community assembly may explain the remarkable evolutionary stasis and long-term persistence of certain microbial lineages in the deep subseafloor sedimentary biosphere.

The finding of this study indicate that the genome structure and gene content of the dominant microbial lineages remained remarkably stable during burial. Across a depth of 31 horizons (∼300 m) and a temporal range of ∼480 kyr, the mutation rates and pN/pS ratios remained constant, showing no systematic increase (Fig. 6). Likewise, gene similarity showed no cumulative decline with depth (Supplementary Fig. 4). These results support the occurrence of evolutionary stasis, whereby microbial populations in energy-limited sediments largely retain genomic signatures established at the time of burial. This interpretation is consistent with earlier observations of genomic stability in shallow sediments^19^ and with the evolutionary stasis reported for deep subsurface lineages such as *Candidatus D. audaxviator*^23^. In contrast, studies on Myr-old isolates from abyssal clays^20^ have reported elevated pN/pS ratios and pseudogene accumulation, emphasising that the evolutionary dynamics in the deep subseafloor are governed less by ongoing mutation and selection than by long-term genomic preservation and ecological stability.

Previous studies have provided compelling evidence of evolutionary stasis in subseafloor microorganisms, particularly in shallow sedimentary settings. For example, Starnawski et al.^19^ suggested evolutionary stasis in Atribacterota and *Dehalococcoidia* within shallow sediments (∼2 mbsf, ∼5 kyr), whereas Garber et al.^21^ showed that *Thalassospira* isolated from 3–6 Myr-old sediments had remained essentially unchanged during burial. Although these studies provided intriguing proof-of-concept evidence, they address either a narrow temporal window in shallow sediments or single cultured lineages. In contrast, our analysis integrated amplicon-based community surveys, genome-resolved metagenomics, and comparative evolutionary metrics (pN/pS, r/m, and genome size) across 31 consecutive horizons spanning up to ∼480 kyr and encompassing multiple phyla. This extensive temporospatial coverage demonstrates that evolutionary stasis is not restricted to specific taxa or exceptional depositional settings but instead represents a general principle of microbial persistence in deep and old subseafloor sediments. Our depth-resolved analyses of the pN/pS ratios (Fig. 6) showed consistently low values across horizons, with no evidence of cumulative increases in non-synonymous variation. The pN/pS ratio is a widely used indicator of selective pressure on protein-coding genes, where low values reflect pervasive purifying selection, which removes deleterious mutations from the population^32,33^. Therefore, the consistently low pN/pS ratios throughout the sediment column indicate that purifying selection predominates across depths, limiting the fixation of amino-acid–altering mutations, even over geological timescales. These results reinforce the idea that genome-wide purifying selection predominates in deep subseafloor sediments, further supporting long-term evolutionary stasis rather than adaptive changes during progressive burial. These findings advance previous work by showing that evolutionary stasis is not restricted to specific taxa or exceptional depositional environments but represents a general pattern of microbial persistence across multiple lineages and depths in the deep subseafloor sedimentary biosphere.

Genomic information obtained from the sediment column at Site C9001 also provides insights into how genome size and coding density vary with sediment depth. Contrary to the expectations of progressive genome streamlining under energy limitation^34,35^, our data showed no evidence of a systematic genome size reduction across the community (Fig. 5). The average genome sizes estimated using MicrobeCensus remained stable throughout the sediment column, and lineage-specific analyses of the major MAG groups showed no clear trends in either genome size or coding density. These results indicate that genome streamlining did not occur during long-term burial. Previous studies have reported that small-genome lineages were preferentially selected during the initial burial process^36^. Consistent with this, our results support the idea that these pre-adapted lineages subsequently persisted under extreme energy limitations, maintaining evolutionary stasis over extended geological timescales. Thus, genome conservation rather than progressive reduction appears to be the dominant principle of microbial persistence in deep subseafloor sedimentary environments.

Another important aspect to consider is the depth-dependent distribution of CAZyme genes, which are central to the degradation and remineralisation of organic carbon, and thus to biogeochemical carbon cycling in marine sediments. No significant differences were observed among MAGs affiliated with Atribacterota JS1 and the Chloroflexota GIF9 clade. This finding indicates that these lineages maintained stable repertoires of genes involved in carbohydrate metabolism throughout the sediment column. Therefore, the absence of depth-dependent changes in CAZyme gene composition among Atribacterota likely reflects a stable, energy-limited environment in which uniform selection maintains functionally conserved metabolic repertoires over geological timescales. Although Dehalococcoidia MAGs exhibited significant shifts in glycoside hydrolase and carbohydrate esterase densities at intermediate depths, these variations lacked a consistent depth-dependent pattern and were not observed in any other lineages (Fig.7, Supplementary Figs. 5 and 6). Such patterns may instead represent the imprint of distinct surface-originating populations isolated in the sediment layer under conditions of dispersal limitation. Overall, the preservation of CAZyme repertoires across depths supports the view that core metabolic traits in Chloroflexota and other lineages primarily reflect legacy effects established prior to burial.

Finally, our analyses of r/m ratios provided insights into the sources of genomic diversity. ClonalFrameML-based estimates indicate that homologous recombination has contributed to genome evolution in several lineages, including Atribacterota, Planctomycetota, and WOR-3 (Table 1). These values are modest compared to highly recombinogenic surface taxa^37^ but exceed the extremely low r/m ratios reported for *Thalassospira* isolates from Myr-old abyssal clays^20^ and for *S. commune* isolates recovered from > 2 km-deep anoxic sediments^22^. Because the r/m estimation from MAGs can, in principle, be inflated by unresolved microdiversity or assembly artifacts, one might question the validity of these values. However, the reference genomes used for comparison in the Genome Taxonomy Database (GTDB) are also MAG-derived; thus, any systematic bias in r/m estimation would be applied uniformly across lineages and would, therefore, unlikely to affect the overall comparative trends reported here. A recent study used MAGs to estimate r/m in natural populations^38^, suggesting that the MAG-based inference of recombination has become an accepted practice in comparative microbial genomics. Based on this evidence, it is likely that most recombination events occurred in surface-to-shallow sediments prior to burial, when cells were more metabolically active, and the population densities were higher. This ecological interpretation of the deep subseafloor sedimentary biosphere is further supported by previous observations of mobile genetic elements such as CRISPR loci and prophages in deep subsurface genomes, which represent a genomic legacy of earlier ecological interactions^23^. Thus, while recombination-derived diversity contributed to shaping subseafloor genomes, subsequent evolution appeared to be dominated by stasis, purifying selection, and the long-term preservation of preexisting variation.

In conclusion, our integrative, depth-resolved analyses demonstrate that microbial genomes in deep subseafloor sediments are largely evolutionarily static, with essential functions maintained under strong purifying selection for hundreds of thousands of years. By spanning 31 consecutive horizons down to ∼300 mbsf and combining community-level assembly processes with genome-resolved evolutionary metrics, our study provides the most comprehensive evidence to date that evolutionary stasis is not an isolated phenomenon but a general principle of microbial persistence in deep subseafloor sedimentary environments. This long-term genomic conservation coexists with limited signals of past recombination and the stable retention of functional repertoires such as CAZymes, highlighting the legacy effects of surface-derived populations. Together, these findings establish a unified perspective on how community assembly and genome evolution interact to shape microbial life under extreme energy limitation, offering a new basis for understanding the resilience and long-term survival of deep subseafloor sedimentary biospheres.

## Methods

### Sediment samples

Sediment samples were collected in 2006 from Site C9001 (41°10.5983′N, 142°12.0328′E; 1180 m water depth) off the Shimokita Peninsula by the D/V Chikyu (Fig. 8) and immediately frozen at −80 °C. In total, 31 sediment core samples were taken at approximately 10 m intervals between 5.2 and 296 mbsf. The sediments are predominantly silty clay, deposited at a rate of 62 cm kyr⁻¹, with an estimated age of ∼480 kyr at the deepest interval^39^. A geothermal gradient of 24 °C km⁻¹ yields in situ temperatures of ∼4 °C near the surface and ∼12 °C at 296 mbsf^1^. Porewater sulphate was already depleted in the uppermost sediments, remained near zero down to ∼100 mbsf, and increased to 0–7 mM below that depth^40^. Microbial abundance, determined by SYBR Green staining, decreased logarithmically with depth from ∼5 × 10⁸ cells cm⁻³ sediment in the shallowest sample to ∼1 × 10⁷ cells cm⁻³ at 200–300 mbsf^1^.

**Fig. 8.**
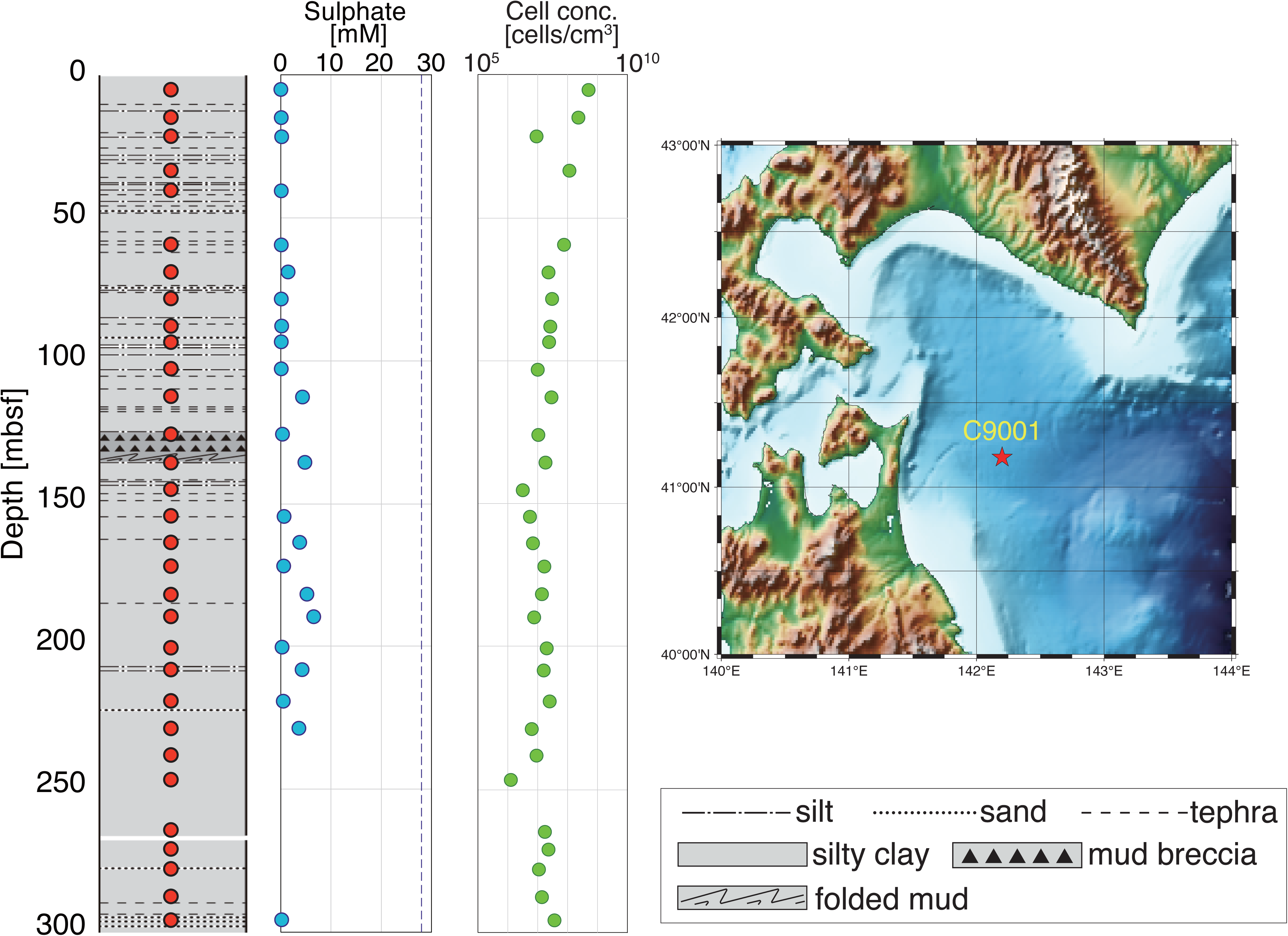
Sediment core from site C9001. Left: Lithology of the core (adapted from^39^). Red circles indicate the depths of the samples used in this study. Middle: Sulphate concentration in the pore water of the sediment samples. Concentration data have been reported previously^40^. The blue dashed line indicates the sulphate concentration of seawater (28 mM). Right: Microbial cell concentration in the sediment samples determined by fluorescence microscopy using SYBR Green I staining^1^. *mbsf* m below the seafloor.

### DNA analysis

DNA was extracted from the frozen sediment using the DNeasy PowerMax Soil Kit (Qiagen) according to the manufacturer’s protocol, except that the kit’s beads were replaced with a 7.5 g + 7.5 g mixture of 0.1 mm and 0.5 mm zirconium beads, and cell disruption was carried out by bead-beating with a Multi-Beads Shocker (Yasui Kikai Co.). The resulting 5 mL DNA eluate was concentrated to 100 µL by ethanol precipitation and further purified using a CHROMA SPIN-1000 column (Takara Bio). For 16S rRNA gene amplicon sequencing, libraries were prepared as described in Hoshino et al.^11^. The V3–V4 region was amplified with the U515F–U806R primer set^41^, and PCR products were purified using NucleoMag NGS Clean-up and Size Select beads (Takara Bio). A second PCR was performed to attach the index and adapter sequences. Amplicons were size-verified by agarose gel electrophoresis, excised, and purified using the NucleoSpin Gel and PCR Clean-up Kit, and sequenced on an Illumina MiSeq system using the MiSeq Reagent Kit v3 (600 cycles). Shotgun metagenomic libraries were prepared using the Nextera DNA Flex Library Prep Kit (Illumina) with 5–20 PCR cycles. Of the 31 libraries generated, 26 were sent to Macrogen Japan for QC and sequencing on the HiSeq X platform, whereas five libraries (2H-6, 5H-4, 12H-1, 22H-6, and 33H-3) were sequenced in-house on a HiSeq 2500, yielding an average of 10.2 Gb of data per sample.

### Data analysis

The amplicon sequences of the 16S rRNA gene were processed as previously described in Hoshino et al.^11^. Briefly, raw reads were quality-trimmed, paired-end merged, and denoised to generate ASVs using USEARCH^42^ (64-bit version). In total, 27 million reads were retained, with an average of 874,000 reads per sample. The taxonomic assignment of each ASV was performed using Mothur^43^ against the SILVA v138 database. Alpha and beta diversity metrics were calculated on datasets rarefied to 200,000 reads per sample using USEARCH and the ‘vegan’ package in R. Community assembly processes were inferred using iCAMP package^27^ in R. As input, 1,275 ASVs, accounting for 95 % of the total reads were selected. A maximum likelihood phylogenetic tree of these ASVs was reconstructed using IQ-TREE^44^.

The ASV tree, abundance matrix, and phylogenetic information were analysed using iCAMP to quantify the relative contributions of the community assembly mechanisms, including homogeneous selection, heterogeneous selection, dispersal limitation, homogenising dispersal, and drift.

### Metagenome Assembly and Binning

Following adapter removal and quality trimming using Platanus_trim v1.0.7, high-quality paired-end reads were assembled de novo using MetaPlatanus^45^. This assembler was selected after benchmarking several tools (including metaSPAdes^46^, MEGAHIT^47^, and IDBA-UD^48^), as it consistently produced assemblies with higher scaffold N50 values, improved contiguity, and better recovery of phylogenetically distinct lineages such as Atribacterota.

For each sample, metagenome-assembled genomes (MAGs) were generated using a combination of MetaBAT2^49^ and MaxBin2^50^. Binning was performed using contigs ≥1500 bp to ensure accuracy, and results from both binning tools were subsequently integrated using DAS Tool^51^ with the default score threshold disabled (--score_threshold 0) to maximise MAG recovery. The completeness and contamination of the resulting MAGs were evaluated with CheckM^52^ using lineage-specific marker gene sets. MAGs with estimated completeness of ≥50% and contamination <10% were classified as medium-or high-quality according to community standards for draft genomes^53^. Taxonomic annotation was performed using several approaches. For each MAG, 16S rRNA gene sequences were identified and compared to the SILVA database (release 138) using BLASTN (e-value ≤1e-5). Whole-genome nucleotide alignments were performed against the NCBI nt and RefSeq databases, while coding sequences were predicted using Prokka^54^ and further annotated by DIAMOND blastp (e-value ≤1e-5) against the NCBI nr database.

Phylogenetic relationships among MAGs were inferred using IQ-TREE^44^. Concatenated alignments of 120 bacterial marker genes and 53 archaeal marker genes, generated by GTDB-Tk^55^, were used as inputs. Maximum likelihood phylogenetic trees were constructed using ModelFinder to select the best-fit substitution model, and branch support was assessed using 1,000 ultrafast bootstrap replicates.

### Gene prediction and functional annotation

Protein-coding sequences were predicted from high-quality MAGs using Prokka. Functional annotation was subsequently performed using MetaCerberus^56^, which integrates multiple curated databases including KEGG^57^, CAZy^58^, MEROPS^59^, Pfam^60^, and TIGRFAMs^61^. MetaCerberus assigns annotations based on HMM-based matching and DIAMOND searches against reference databases, enabling the comprehensive identification of metabolic and functional gene content. Annotations assigned to the CAZy database were extracted from the MetaCerberus output for further analysis of CAZyme profiles.

For quantitative analyses, we selected taxonomic classes containing at least 15 high-quality MAGs (≥90% completeness and <5% contamination), which included Atribacterota JS1 clade, *Dehalococcoidia*, and the Chlofoflexota GIF9 clade. For each MAG, the number of genes assigned to each CAZy category was normalised by genome size (in megabases) to calculate CAZyme gene density. These normalised values were compared across three sediment depth intervals (0–100 m, 100–200 m, and 200–300 mbsf). Statistical significance of depth-related differences was assessed using the two-tailed Mann–Whitney U test, with p <0.05 considered significant.

### Analysis of recombination and mutation rates (r/m)

To assess the overall genomic fluidity and the relative impact of recombination versus mutation on the diversification of key lineages, we performed a population genomic analysis using ClonalFrameML^62^ This method estimates key population genetic parameters from a multiple sequence alignment and the accompanying phylogeny, requiring the construction of a core-genome alignment from all MAGs belonging to a target species-level cluster (ANI>97%). The analysis was applied to clusters containing five or more high-quality genomes, including the MAGs recovered in this study, as well as comparative clusters of high-quality MAGs retrieved from the GTDB^63^ (specifically, those belonging to the phylum Atribacterota class JS1, the Planctomycetota family *Anaerobacaceae*, WOR-3, and class E44-bin80). For each cluster, the required inputs for ClonalFrameML were generated through a robust pipeline. Protein-coding genes and their corresponding nucleotide sequences were predicted for each MAG using Prodigal ^64^. The resulting proteomes were processed using OrthoFinder^65^ to identify a set of single-copy orthologous genes. Subsequently, a supermatrix was constructed using a custom pipeline: for each single-copy ortholog, nucleotide sequences were aligned as codons by using their MAFFT^66^-aligned protein translations as a template in pal2nal.pl^67^. Alignments that failed a validation were automatically discarded before all valid codon alignments were concatenated into a final supermatrix using AMAS^68^. This concatenated alignment was then used to infer a maximum likelihood phylogenetic tree in IQ-TREE^44^ with 1,000 ultrafast bootstrap replicates for branch support. Finally, after unrooting the tree with the R package ‘ape’, we ran ClonalFrameML using the unrooted tree and the supermatrix to estimate key population genetic parameters, including R/θ, 1/δ, and ν.

### Clustering-based analysis of depth-dependent variation in predicted genes

To evaluate depth-related trends in nucleotide variation and selective pressure at the gene level, we focused on bacteria belonging to four phyla (Atribacterota, Chloroflexota, Candidatus Aerophobetes, and Actinomycetota). The predicted gene sequences were obtained using Prodigal from the metagenome-assembled draft genomes reconstructed with Metaplatanus. For each phylum, amino acid sequences of predicted genes from the shallowest sample (depth 1) were compared to those from deeper samples using BLASTp, and sequences sharing ≥97% identity across the full alignment length were selected as candidate orthologues. Candidate genes were clustered using genes detected in the shallowest sample as the references. If multiple candidates were identified, then the sequence with the highest bit score was retained. Subsequently, nucleotide sequences of selected candidates were further compared using BLASTn (≥97% identity across full-length alignments), and the clustering process was validated. Genes for which a 1:1 match to the depth 1 gene could not be confidently determined (i.e., multiple candidates with identical bit scores) were excluded. Additionally, only gene clusters detected in ≥80% of the sampled depths were retained for further analysis. Before clustering, we excluded depths that contained insufficient ortholog candidates and would otherwise decrease the number of gene clusters ultimately recovered. For each gene cluster, the number of synonymous and nonsynonymous substitutions between depth 1 and other depths was counted using the KaKs Calculator^69^, and the variation rate and pN/pS ratios were calculated as described by Starnawski et al.^19^.

## Acknowledgements

The authors express their gratitude to all the crew members of the *Chikyu* shake-down expedition in 2006 (CK06-06), expedition project managers, shipboard scientists, operational staff, and laboratory technicians for their contributions to core sampling and shipboard measurements. We thank the laboratory technicians at Kochi Institute for Core Sample Research, JAMSTEC, for their technical assistance with the experiments. Parts of the computational analyses were facilitated using the Earth Simulator supercomputer (JAMSTEC), and minor assistance in streamlining some bioinformatics coding tasks was provided by ChatGPT (OpenAI). This work was partially supported by the Japan Society for the Promotion of Science (JSPS) Grant-in-Aid for Scientific Research JP22H00429 (to F.I., T.H., and H.T.), JP16H06279, JP1703956, JP21K19876, and JP23K22618 (to T.H.); JST ASPIRE for Top Scientists (to F.I. and T.H.); and JSPS World Premier International Research Center Initiative (WPI: to F.I. and T.H.); Alfred Sloan Foundation Deep Life Community pilot project of the Deep Carbon Observatory (to T.H.); and NINS Astrobiology Center program (AB041005 to T.H.).

## Contributions

T.H. and F.I. conceptualised and designed the study. T.H. supervised the experiments, including the DNA extaction and sequencing. T.H., A.S., R.K., T.I., and H.D. analysed DNA data including amplicon and shotgun metegemic sequencing. T.H. and F.I wrote the manuscript with contributions from all coauthors.

## Data Availability

All sequencing data generated in this study have been deposited in the DDBJ/ENA/NCBI Sequence Read Archive under accession numbers SAMD01688496-SAMD01688527, SAMD00601979-SAMD00601983, and SAMD01591414-SAMD01591445. Additional data are available in the Supplementary Information and Source Data files.

