## Supplementary fig. 1 for "Evolutionary stasis and homogeneous selection structure microbial communities in the deep subseafloor sedimentary biosphere"

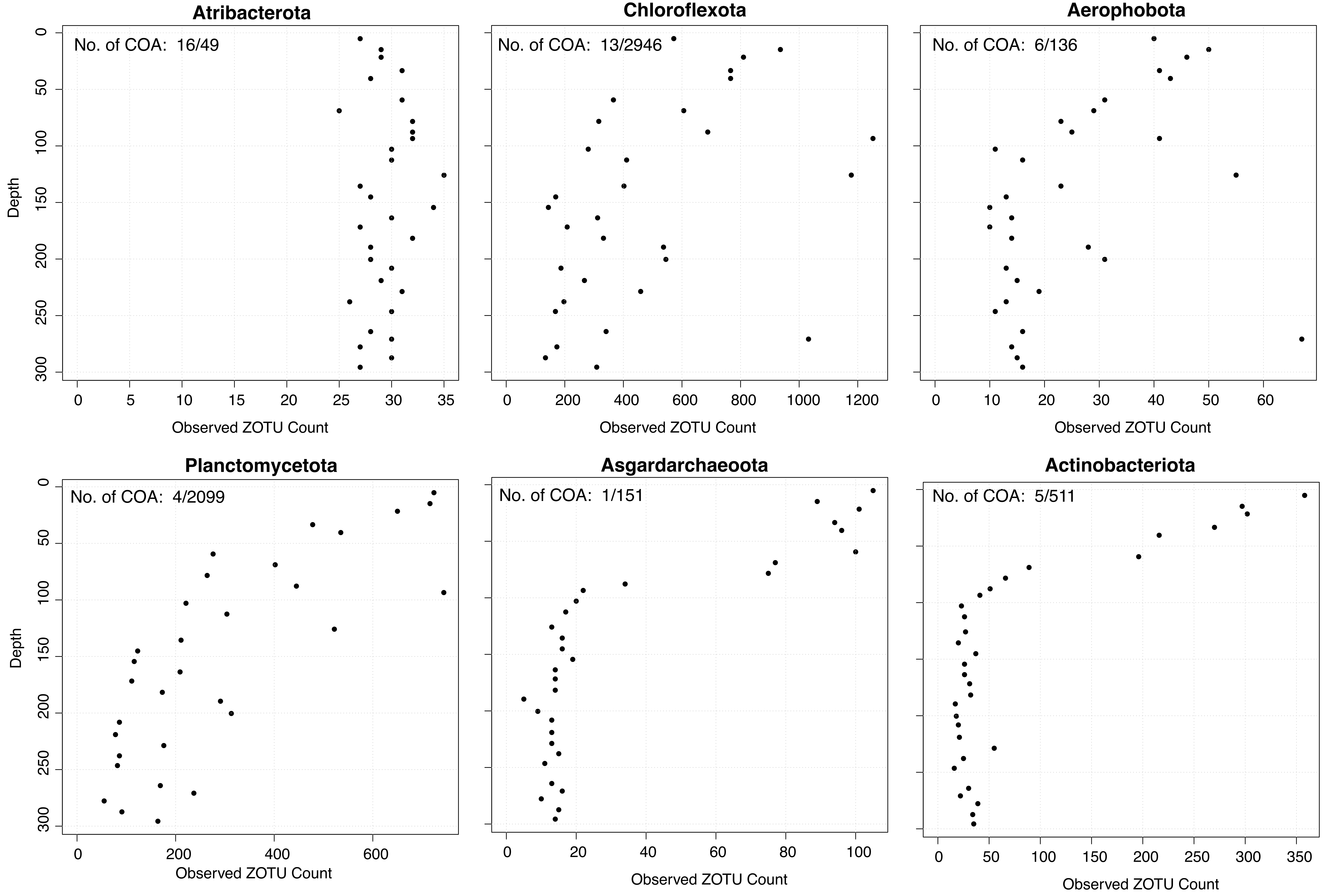


**Supplementary fig. 1 Depth profile of observed ASV (Amplicon Sequence Variant) counts.**
The ASV counts were standardized by rarefying each sample to a depth of 200,000 reads, allowing for consistent comparisons of microbial diversity across different depths. COA indicates consistently observed ASVs across the depth profile. “No. of COA” shows, for each phylum, the number of consistently observed ASVs relative to the total number of ASVs detected in that phylum (COA / total).
