## Supplementary fig. 2 for "Evolutionary stasis and homogeneous selection structure microbial communities in the deep subseafloor sedimentary biosphere"

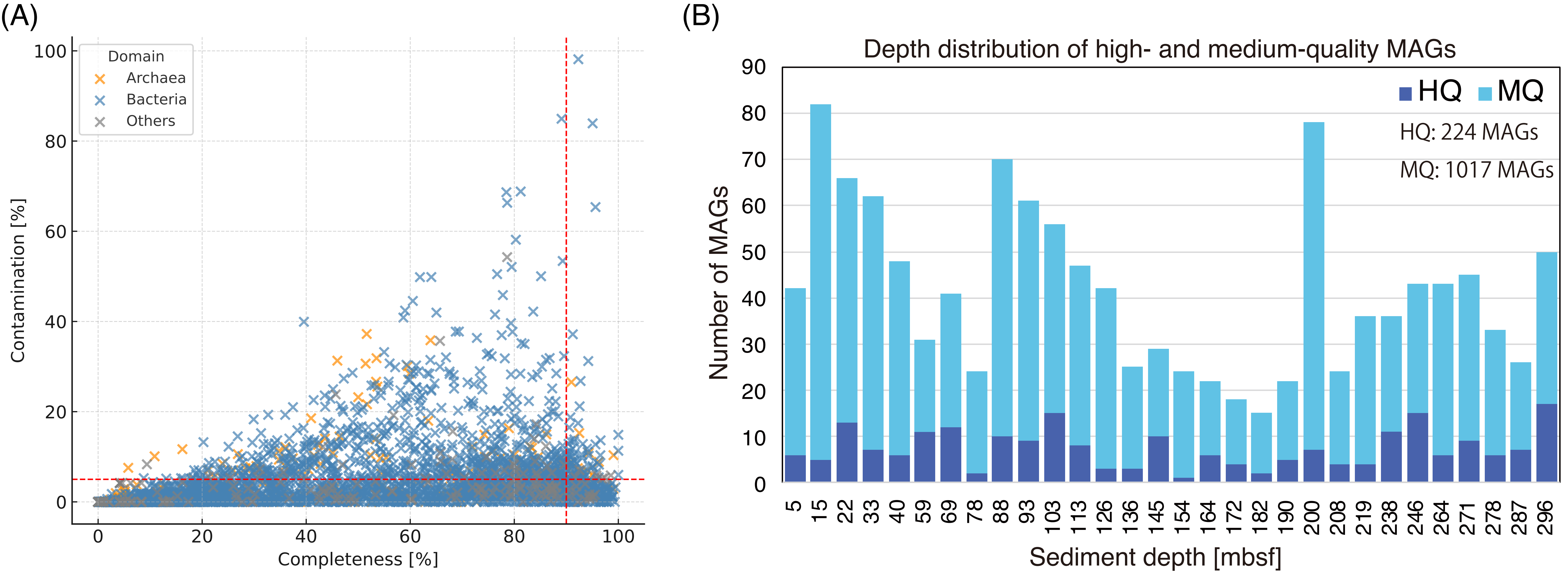


**Supplementary fig. 2:**

(A) Scatter plot of 3,993 genome bins obtained from subseafloor sediment. Colors represent taxonomic domains: Bacteria (blue), Archaea (orange), and Others (gray). Among these bins, 224 were classified as high-quality metagenome-assembled genomes (HQ MAGs) based on the criteria of >90% completeness and <5% contamination. Red dashed lines indicate these thresholds. (B)Bar plot showing the number of metagenome-assembled genomes (MAGs) recovered from individual sediment depths.

MAGs are classified as high-quality (HQ: dark blue; completeness > 90%, contamination < 5%) or medium-quality (MQ: light blue; completeness 50–90%, contamination < 10%).
