## Supplementary fig. 3 for "Evolutionary stasis and homogeneous selection structure microbial communities in the deep subseafloor sedimentary biosphere"

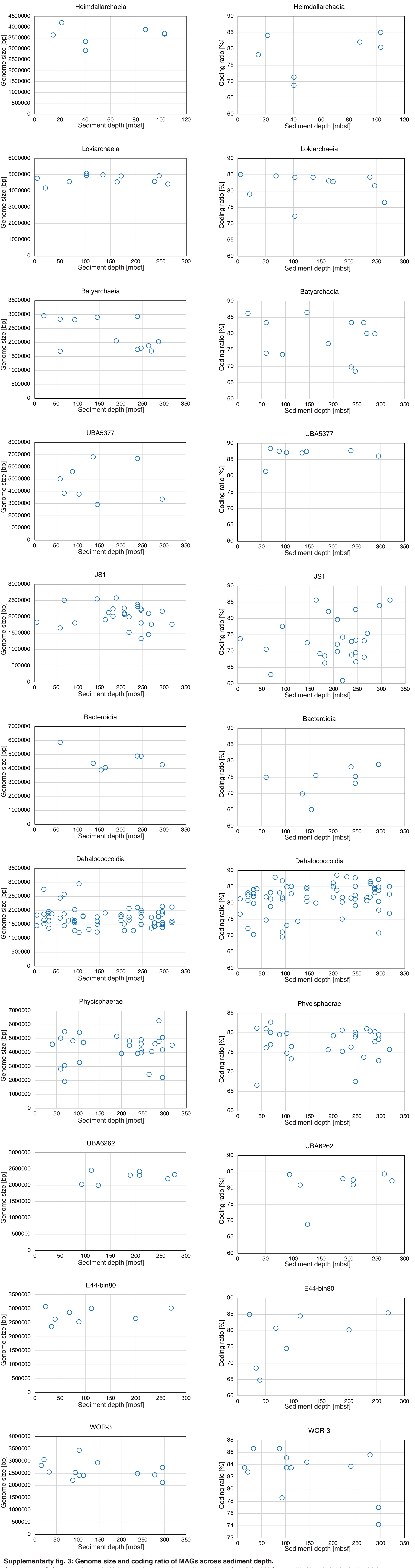


**Supplementarty fig. 3: Genome size and coding ratio of MAGs across sediment depth.**
Genome size (left) and coding ratio (right) are plotted against sediment depth (mbsf) for MAGs classified into individual microbial classes. No consistent depth-related trend is observed within each lineage.
