## Supplementary fig. 4 for "Evolutionary stasis and homogeneous selection structure microbial communities in the deep subseafloor sedimentary biosphere"

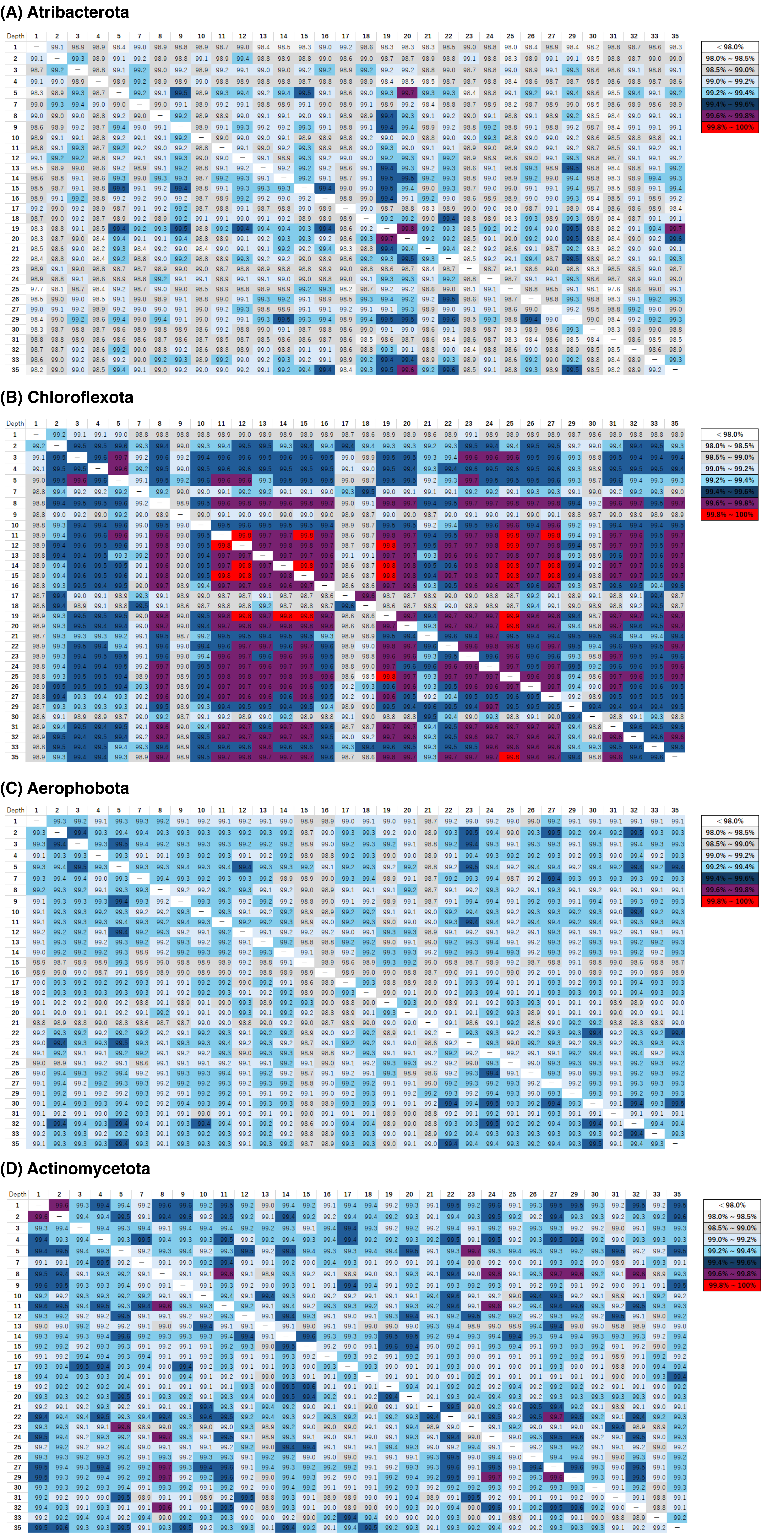


**Supplementary fig. 4: Sequence similarity of orthologous genes across sediment depths.**
For four bacterial lineages (Atribacterota, Chloroflexota, Aerophobota and Actinomycetota), predicted genes from metagenome-assembled genomes were compared across depths by all-vs-all BLASTn. Only genes with ≥97% nucleotide identity across full-length alignments were retained, and the average pairwise sequence identity between depths was calculated. Across all lineages examined, sequence similarity remained uniformly high, with no systematic decline as depth increased. These results indicate that nucleotide substitutions do not accumulate progressively with burial depth, supporting the view that genetic variation remains largely static in subseafloor microbial populations.
