## Supplementary fig. 5 for "Evolutionary stasis and homogeneous selection structure microbial communities in the deep subseafloor sedimentary biosphere"

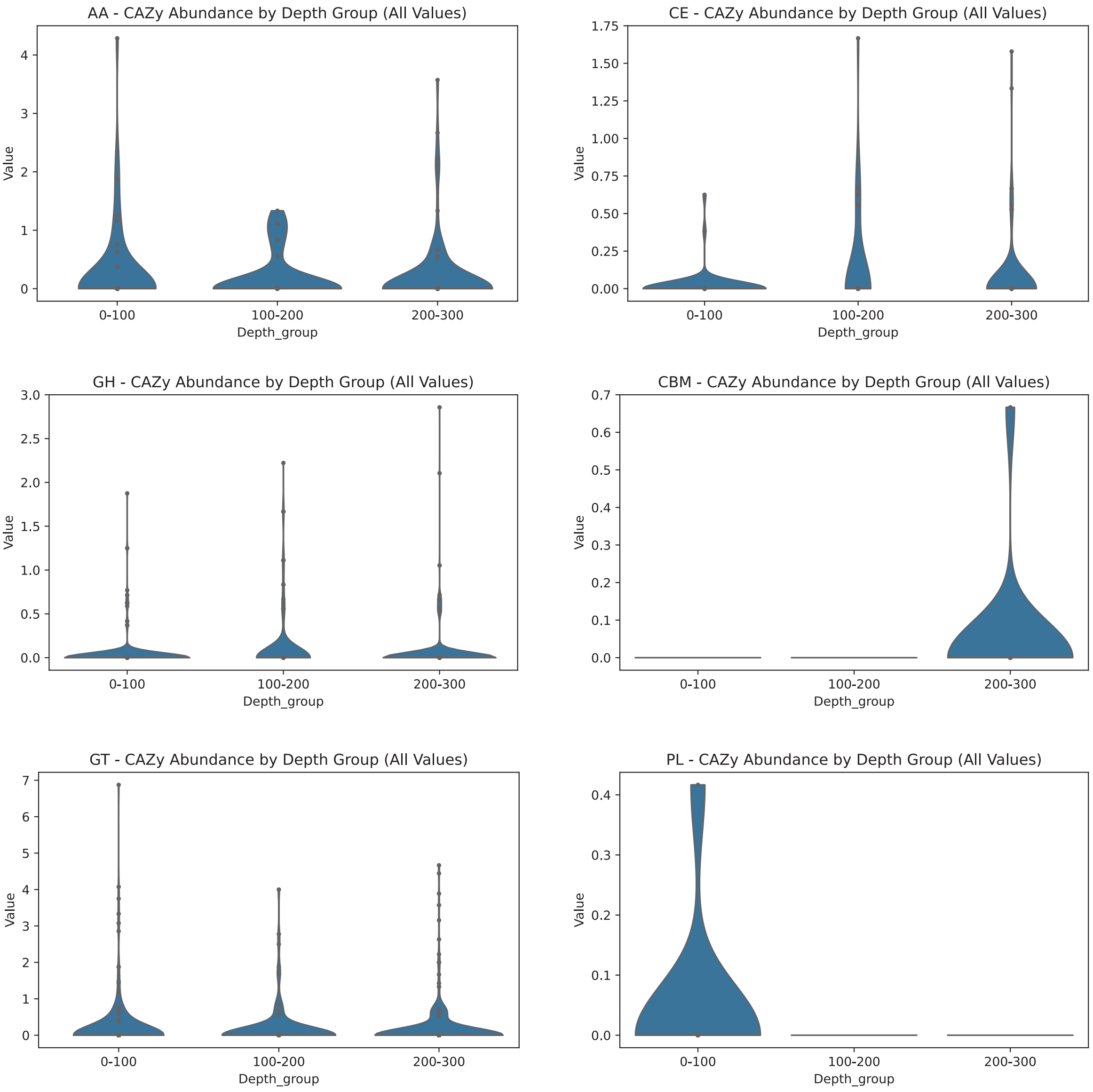


| Category | Group1 | Group2 | U_statistic | p_value | n_Group1 | n_Group2 |
|---|---|---|---|---|---|---|
| AA | 0-100 | 100-200 | 1124 | 0.64899061 | 60 | 36 |
| AA | 0-100 | 200-300 | 3074 | 0.30542596 | 60 | 96 |
| AA | 100-200 | 200-300 | 1783 | 0.67351289 | 36 | 96 |
| GH | 0-100 | 100-200 | 36519 | 0.00560248 | 360 | 216 |
| GH | 0-100 | 200-300 | 102264.5 | 0.375686 | 360 | 576 |
| GH | 100-200 | 200-300 | 65204 | 0.02071943 | 216 | 576 |
| GT | 0-100 | 100-200 | 12592 | 0.24657285 | 200 | 120 |
| GT | 0-100 | 200-300 | 32617 | 0.56939917 | 200 | 320 |
| GT | 100-200 | 200-300 | 18642 | 0.44990672 | 120 | 320 |
| CE | 0-100 | 100-200 | 626.5 | 0.01886843 | 50 | 30 |
| CE | 0-100 | 200-300 | 1901 | 0.28409626 | 50 | 80 |
| CE | 100-200 | 200-300 | 1338 | 0.0997292 | 30 | 80 |
| CBM | 0-100 | 100-200 | 30 | 1 | 10 | 6 |
| CBM | 0-100 | 200-300 | 75 | 0.47676672 | 10 | 16 |
| CBM | 100-200 | 200-300 | 45 | 0.60983404 | 6 | 16 |
| PL | 0-100 | 100-200 | 33 | 0.51860502 | 10 | 6 |
| PL | 0-100 | 200-300 | 88 | 0.23567991 | 10 | 16 |
| PL | 100-200 | 200-300 | 48 | 1 | 6 | 16 |

**Supplementary fig. 5: Depth-dependent variation in CAZyme gene density across MAGs affiliated with Dehalococcoidia.**

Violin plots depict the normalized gene densities of each CAZyme category across three depth intervals. Significant differences were observed in glycoside hydrolase (GH) densities between 0–100 vs 100–200 m ($p = 0.0056$) and 100–200 vs 200–300 m ($p = 0.021$), and in carbohydrate esterase (CE) densities between 0–100 and 100–200 m ($p = 0.0189$). Other CAZyme categories showed no significant depth-related trends.
