## Supplementary fig. 6 for "Evolutionary stasis and homogeneous selection structure microbial communities in the deep subseafloor sedimentary biosphere"

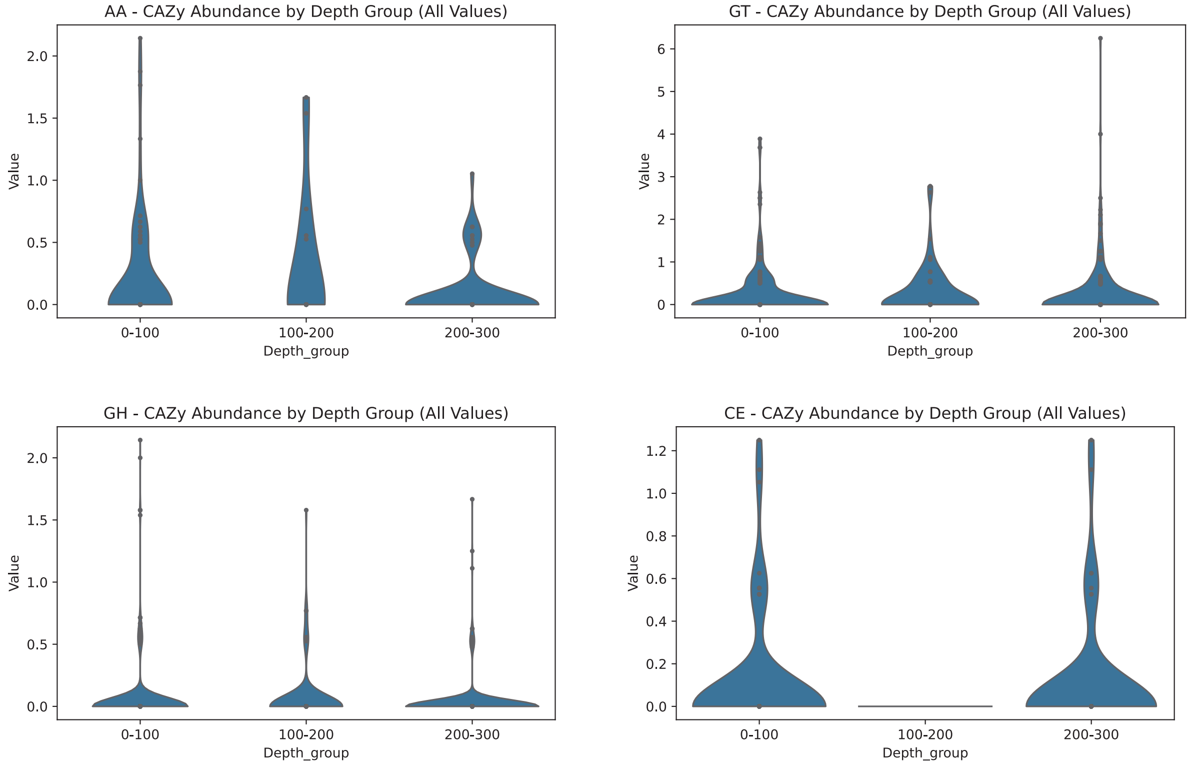


| Category | Group1 | Group2 | U_statistic | p_value | n_Group1 | n_Group2 |
|---|---|---|---|---|---|---|
| AA | 0-100 | 100-200 | 641 | 0.46769461 | 78 | 18 |
| AA | 0-100 | 200-300 | 2604 | 0.11196185 | 78 | 60 |
| AA | 100-200 | 200-300 | 647 | 0.06628297 | 18 | 60 |
| GH | 0-100 | 100-200 | 24760.5 | 0.47662123 | 468 | 108 |
| GH | 0-100 | 200-300 | 86000 | 0.22412689 | 468 | 360 |
| GH | 100-200 | 200-300 | 20239 | 0.12109815 | 108 | 360 |
| GT | 0-100 | 100-200 | 7287.5 | 0.29476683 | 260 | 60 |
| GT | 0-100 | 200-300 | 26578.5 | 0.57515227 | 260 | 200 |
| GT | 100-200 | 200-300 | 6515.5 | 0.17197475 | 60 | 200 |
| CE | 0-100 | 100-200 | 562.5 | 0.10991401 | 65 | 15 |
| CE | 0-100 | 200-300 | 1644 | 0.86572704 | 65 | 50 |
| CE | 100-200 | 200-300 | 322.5 | 0.13240755 | 15 | 50 |

**Supplementary fig. 6: Depth-dependent variation in CAZyme gene density across MAGs affiliated with the GIF9 clade.**

Normalized CAZyme gene densities are shown across three sediment depth intervals. No statistically significant differences were detected across depths for any CAZy category ($p > 0.05$), indicating functional stability in carbohydrate metabolism among GIF9 MAGs throughout the sediment column.
