## Supplementary fig. 7 for "Evolutionary stasis and homogeneous selection structure microbial communities in the deep subseafloor sedimentary biosphere"

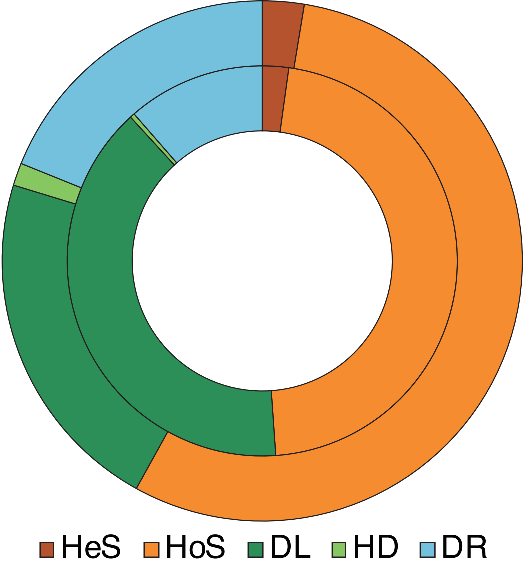


**Supplementary fig. 7: Community assembly mechanisms of microbial communities in shallow and deep marine sediments.** The relative contributions of different ecological processes were quantified using iCAMP. The inner circle represents shallow sediments (<100 mbsf), while the outer circle represents deep sediments (>100 mbsf).

HeS, heterogeneous selection; HoS, homogeneous selection; DL, dispersal limitation; HD, homogenizing dispersal; DR, drift.
