## Supplementary Table 1 for "Evolutionary stasis and homogeneous selection structure microbial communities in the deep subseafloor sedimentary biosphere"

**Supplementary Table 1. Taxonomic summary of ASVs showing significant increases (A) or decreases (B) in relative abundance with depth.**

(A)

| Phylum | No. of ASVs | Abundance [%] |
|---|---|---|
| Actinobacteriota | 3 | 0.603 |
| Chloroflexota | 2 | 3.102 |
| Deinococcota | 2 | 0.015 |
| Proteobacteria | 2 | 0.171 |
| Bacteroidota | 1 | 0.127 |
| Aerophobota | 1 | 11.831 |
| Firmicutes | 1 | 0.001 |

(B)

| Phylum | No. of ASVs | Abundance [%] |
|---|---|---|
| Actinobacteriota | 250 | 3.875 |
| Chloroflexota | 245 | 3.798 |
| Planctomycetota | 204 | 3.162 |
| Bacteria_unclassified | 201 | 3.116 |
| Thermoplasmatota | 138 | 2.139 |
| Firmicutes | 123 | 1.907 |
| Proteobacteria | 119 | 1.845 |
| Asgardarchaeota | 94 | 1.457 |
| Crenarchaeota | 75 | 1.163 |
| Armatimonadota | 71 | 1.101 |
| Nanoarchaeota | 68 | 1.054 |
| Desulfobacterota | 62 | 0.961 |
| Archaea_unclassified | 44 | 0.682 |
| Acidobacteriota | 40 | 0.620 |
| Spirochaetota | 32 | 0.496 |
| Bacteroidota | 32 | 0.496 |
| Patescibacteria | 25 | 0.388 |
| Aerophobota | 18 | 0.279 |
| Verrucomicrobiota | 9 | 0.140 |
| Marinimicrobia_(SAR406_clade) | 7 | 0.109 |
| WS1 | 7 | 0.109 |
| Schekmanbacteria | 6 | 0.093 |
| Acetothermia | 6 | 0.093 |
| Sva0485 | 6 | 0.093 |
| WS4 | 6 | 0.093 |
| Calditrichota | 5 | 0.078 |
| Cyanobacteria | 5 | 0.078 |
| DTB120 | 5 | 0.078 |
| Euryarchaeota | 5 | 0.078 |
| Zixibacteria | 5 | 0.078 |
| Myxococcota | 4 | 0.062 |
| Elusimicrobiota | 4 | 0.062 |
| BHI80-139 | 3 | 0.047 |
| WS2 | 3 | 0.047 |
| TA06 | 3 | 0.047 |
| Halobacterota | 2 | 0.031 |
| uncultured | 2 | 0.031 |
| Atribacterota | 2 | 0.031 |
| Sumerlaeota | 2 | 0.031 |
| Gemmatimonadota | 2 | 0.031 |
| Latescibacterota | 2 | 0.031 |
| Aenigmarchaeota | 1 | 0.016 |
| Nitrospirota | 1 | 0.016 |
| NB1-j | 1 | 0.016 |
| Campilobacterota | 1 | 0.016 |
| SAR324_clade(Marine_group_B) | 1 | 0.016 |
| Dependentiae | 1 | 0.016 |

ASVs showing a significant positive or negative correlation with sediment depth were identified based on Spearman's rank correlation coefficient ($\rho \geq 0.5$ or $\rho \leq -0.5$, $p < 0.05$). The table summarizes the number of such ASVs per phylum and their total relative abundance (%) across all samples.
